# Distinct genetic signatures of cortical and subcortical regions associated with human memory

**DOI:** 10.1101/524116

**Authors:** Pin Kwang Tan, Egor Ananyev, Po-Jang (Brown) Hsieh

## Abstract

Despite the discovery of gene variants linked to memory performance, understanding the genetic basis of human memory remains a challenge. Here, we devised a framework combining human transcriptome data and a functional neuroimaging map to uncover the genetic signatures of memory in functionally-defined cortical and subcortical memory regions. Results were validated with animal literature and our framework proved to be highly effective and specific to the targeted cognitive function versus a control function. Genes preferentially expressed in cortical memory regions are linked to associative learning and ribosome biogenesis. Genes expressed in subcortical memory regions are associated with synaptic signaling and epigenetic processes. Cortical and subcortical regions share a number of memory-related biological processes and genes, e.g. translational initiation and *GRIN1*. Thus, cortical and subcortical memory regions exhibit distinct genetic signatures that potentially reflect functional differences in health and disease, and propose gene candidates for the targeted treatment of memory disorders.

## INTRODUCTION

Memory function is crucial for everyday life. It is involved in a wide variety of cognitive tasks, from mental arithmetic to long-term planning. Human memory function is relatively well characterized in terms of gross neural correlates related to behavior and mental disorders. Insights gained from non-invasive functional magnetic resonance imaging (fMRI) and brain lesion case studies^1–3^ led to an understanding of cortical and subcortical memory regions as functionally distinct areas subsumed under the broad umbrella of memory function. Yet, despite the fact that memory ability is highly heritable^4,5^, with genetic risk factors for memory disorders^6^, the genetic signature underlying human memory remains poorly understood^7^. This gap can potentially be addressed by combining recently created human brain transcriptomes^8^ and neuroimaging maps^9–11^. While similar work has been done in recent years, most effort concentrate either on functional networks in either the cerebral cortex or subcortex due to the disparate patterns of gene expression across these regions^8,12–16^. However, recent work also suggests that functional networks in the human brain have convergent gene expression patterns across cortical and subcortical areas^17^. Therefore, a key question stands: Do functionally distinct human cortical and subcortical memory regions^18–21^ possess distinct genetic signatures associated with memory function? The answer could provide an insight into the underlying biological processes and genes underlying human cortical and subcortical memory, and benefit drug discovery for cognitive enhancement^22^ and the treatment of memory disorders^23–25^.

Exploring the genetic basis of cortical and subcortical memory is relevant to understanding the distinction between different memory types, stages of memory formation, and the associated cognitive functions and disorders. Human memory is broadly classified into declarative and non-declarative memory^26^, and has been studied with fMRI and lesion studies. Explicit or declarative memory involves conscious awareness and is generally associated with facts, events, and places. Implicit or non-declarative memory does not require conscious awareness and is linked to perceptual and motor learning^7^. Functionally, explicit memory is largely linked to activity in cortical regions (hippocampus and perirhinal, entorhinal, and parahippocampal cortices), while implicit memory is related to subcortical regions (amygdala, striatum, basal ganglia cerebellum)^7,20,21,27^. This divide in cortical-subcortical function is also important in various facets of healthy memory function and memory disorders. In systems consolidation, memory consolidation is primarily associated with cortical regions, including the hippocampus^28^. Among cognitive disorders, Alzheimer’s disease^29^ and schizophrenia^30–32^ are also associated with cortical mechanisms. In contrast, memory disorders such as dementia may be classified as either cortical or subcortical, depending on behavioral and neuropsychological deficits observed ^18,19,33^. As such, the genetic signature of cortical and subcortical memory may yield insight into the various memory processes and disorders specifically associated with each region. Additionally, understanding the general genetic mechanisms supporting human memory function is crucial given the role of memory in development^34^, ageing^35–37^, and numerous psychiatric disorders. The latter include undesirable enhanced memory in post-traumatic stress disorder and impaired modulation between memory and reward systems in addiction^38^. Furthermore, identifying the genetic signatures of human memory may be useful for drawing comparisons between the molecular mechanisms of memory in models of animal and human cognition^39,40^. Importantly, a drug discovery approach centered on human genomics^41–43^ that is complemented with animal research may accelerate current efforts in identifying drug candidates for memory disorders^44–46^.

Given that diverse memory processes have common cortical or subcortical neural substrates, we sought to establish if there are unifying genetic signatures across cortical or subcortical memory areas. Convergent gene expression across cortical or subcortical memory regions may be expected due to anatomical and/or functional relatedness in various stages of life. During neurodevelopment, functionally-related neurons may migrate across cortical or subcortical areas, and thus form a convergent gene expression profile across these regions^47,48^. During adulthood, neurons in different brain areas may be structurally^49,50^ or functionally connected^51^, and thus share a common molecular mechanisms for areas within the same functional network^12–14,17,52,53^. These shared gene signatures may reflect cortico-cortical^13^ and functional connectivity^14,54^. Such trends may also extend to memory, as such functional and structural connectivity is also observed seen between cortical areas linked to memory performance. Functional networks underlie memory performance in the cortex^55,56^, including a cortex-based medial-temporal-lobe-and-neocortex network involved in episodic memory and memory encoding^57–60^. Structural connectivity in cortical areas is also tied to episodic memory performance in children and adults^61,62^. Given that spatial gene expression is known to underlie functionally and structurally connected areas, we may expect to find gene signatures of cortical and subcortical areas linked to memory. However, these cortical and subcortical gene signatures are likely distinct due to their roles in health and disease.

Here we developed a novel, validated framework that identifies gene signatures of human cortical and subcortical memory. We first correlated the spatial human brain transcriptomes and neuroimaging data for each gene^15^ (**Fig. 1a, b**). Second, we analyzed both cortical and subcortical regions separately, and identified biological processes and candidate genes with Gene Set Enrichment Analysis (GSEA) and Leading edge analysis (LEA) respectively (**Fig. 1c, d**). GSEA analyzes genes as potentially working towards a common biological function, instead of independent individual entities^63,64^. GSEA and LEA were effective in identifying genetic signatures of cognitive functions^65–67^, including episodic and working memory^68,69^. Third, we validate the link between these discovered genes and human memory by drawing from animal memory literature (**Fig. 1e**). Fourth, we assess method effectiveness and specificity in identifying genes strictly related to memory function as opposed to a control function i.e. double dissociation (**Fig. 1f**), and vice versa. With this framework, we found largely distinct memory-related biological processes and genes across cortical and subcortical regions. These results indicate that the cortical and subcortical memory regions have largely distinct genetic signatures. These biological processes and genes may provide a better understanding of genetics that underlie systems-level memory, and a potential basis for the targeted treatment of memory disorders^21^.

**Figure 1.**
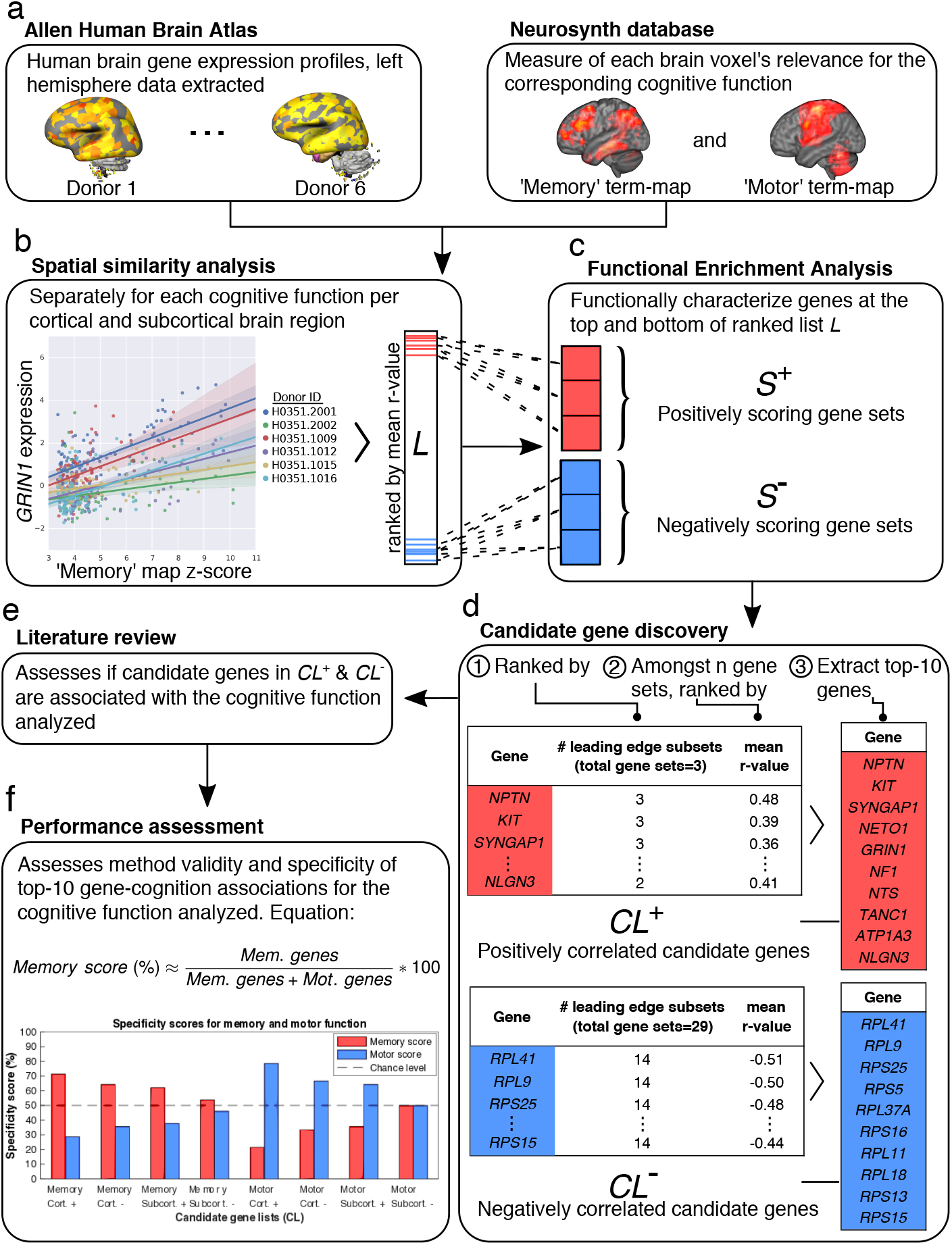
Overview of genetic signature discovery framework. **a**, Preprocessing of Allen Human Brain Atlas and Neurosynth neuroimaging maps, and their integration into common neuroimaging template space. **b**, Calculation of spatial similarity between the maps separately for the memory and motor functions, and for the cortical and subcortical regions, giving four ranked gene lists *L* in total (contains genes and mean *r*-value). **c**, Functional characterization of each *L* with biologically meaningful gene sets with GSEA-Preranked analysis (dotted lines connecting *L* and gene sets represent the clustering of genes into enriched gene sets), yielding positively and negatively scoring gene sets *S*^+^ and *S*^−^. **d**, Identification of candidate genes associated with the cognitive function and brain region, operationalized as the subset of genes driving the enrichment score of the significantly enriched gene sets found using GSEA-Preranked analysis. This produced two candidate gene lists, *CL*^+^ and *CL*^−^, containing highly positively and negatively correlated genes from *S*^+^ and *S*^−^, respectively. **e**, Literature review of each *CL* quantifying the genes associated with the target or control cognitive function. **f**, Assessing framework validity and specificity with each of eight *CL*s. (2-column image, colored)

## MATERIALS AND METHODS

### Allen Human Brain Atlas

The AHBA transcriptome was generated from normalized mRNA microarray sampling of a combined 3702 sampling sites across six donor brains aged 24 to 57 (*N* = 6 left hemispheres, *N* = 2 right hemispheres)^8^. Data from all six donors was horizontally concatenated into a .csv file, with one probe per row.

### Neurosynth memory association map

Neurosynth association maps (thresholded FDR < 0.01) were used as neuroimaging data for the memory and motor functions^9^. These cognitive functions were chosen as they were largely functionally and anatomically distinct, constructed from a similar number of studies (N_memory_ = 2744, N_motor_ = 2565). Neurosynth quantifies the relevance of each voxel to the user-specified search terms based on a database of neuroimaging studies. Each voxel is assigned a z-score reflecting the number of studies with the search terms in the title or the abstract. Negative z-scores indicate a higher correlation with other search terms unrelated to memory, and thus were excluded from our analyses. For broad cognitive function domains, single terms enable the generation of maps that approximate the target cognitive process reasonably well^9^. Therefore, we used ‘memory’ and ‘motor’ as our search terms to derive the memory and motor association maps. Note that in the example of memory, this approach resulted in inclusion of a broad range of sub-functions, such as working memory and long-term memory. This allowed for a broader definition of memory and motor function for subsequent candidate gene identification steps.

### AHBA mRNA probe election procedure

To accurately estimate wild-type gene expression, probes were collapsed with a one-probe-per-gene representation as per Richiardi *et al*. (2015)^52,70^ instead of averaging across probes^15^. Genes without an official Entrez ID were excluded. We also excluded the probes that were either expressed insignificantly above background, or were not mapped to the gene’s exonic region. For each remaining gene, the probe with the best match to the gene’s exonic region was selected^52^. This probe selection procedure was meant to account for the varying reliability of AHBA probe measurements due to microarray probe-gene mismatches, which could lead to a misrepresentation of true gene expression level^70^.

### Defining cortical and subcortical regions

The data were divided into cortical and subcortical regions with the modified Brodmann atlas^8^ used by AHBA. The AHBA data was normalized (z-score) for all samples per donor, separately for cortex (which includes the hippocampus) and subcortex, with the z-score indicating the expression level in normalized units. This was done to account for individual variation in gene expression^71^. This gave an individual average of 242 left cortical (range: 196 to 287) and 201 left subcortical samples/transcriptomes per donor (range: 158 to 232) before further restriction to brain regions of interest.

### Co-registration of AHBA and Neurosynth memory map

To allow a comparison of spatial similarity between neuroimaging and AHBA maps of differing resolutions, both maps were transformed into a common 3D stereotactic brain space. This was done by mapping AHBA data point coordinates to the Montreal Neurological Institute 152 (MNI152) template space. The Neurosynth map was used as a mask for the AHBA map, so that only the overlapping areas were included in the correlation analysis. Due to the limited availability of hemispheres sampled (six left and two right hemispheres), we used only the left hemispheres, separated into cortical and subcortical regions. In subsequent steps, the cortical and subcortical analyses were kept separate. In addition to allowing insight into shared and dissociated cortical/subcortical genetic mechanisms, this also avoided confounds from the divergent transcriptional profiles^52^. We then matched the smoothing of both maps by smoothing the AHBA with a 6mm radius sphere^15^. At the end of this step, there remained on average 117 (range: 85 to 138) memory and 71 (range: 60 to 82) motor cortical usable data points, and on average 47 (range: 39 to 56) memory and 103 (range: 84 to 131) motor subcortical usable data points per individual.

### Spatial correlation analysis of AHBA and Neurosynth data

Fox *et al*. (2014)^15^ developed a tool for correlating the spatial AHBA and neuroimaging maps^9^ to identify top correlated genes as candidate for gene-cognition associations^15^. Their method leveraged on findings that the reward-associated dopamine receptor gene (*DRD2*) was consistently and highly expressed in the striatum^72–74^, a brain area reliably found to be relevant for reward-processing^75,76^. This suggests that the spatial intensity of genetic expression signatures may correlate with the neural correlates of specific cognitive functions. We used the spatial analysis separately for cortical and subcortical regions. An approximate random effects analysis was used to account for individual gene expression variability and to counter the sparse cortical sampling in the AHBA maps^15^. Donor regression slope and intercept were modeled individually. This returned each gene’s mean correlation value (averaged across the six donors), which was the statistic of interest. For a candidate gene associated with memory, we would expect high spatial similarity between both maps, i.e., a pattern of high gene expression within areas highly relevant for memory and vice versa. This would be reflected in a high mean correlation value for that gene. As a result of this step, we obtained four lists *L* of 16906 genes, for memory/motor functions, and cortical/subcortical regions.

### Identifying biological processes of cognitive function

We used a gene set analysis tool (GSEA Pre-ranked) to identify sets of genes associated with common biological functions. The four lists of genes *L* were ranked by mean correlation value and passed to GSEA Pre-ranked. For GSEA Pre-ranked, we analyzed each list *L* with the GSEA Pre-ranked module (v5) on GenePattern with the default parameters^63,77^, including weighted scoring using the Gene Ontology Biological Process library (c5.bp.v6.0.symbols.gmt). GSEA-Preranked looks separately at the top and bottom of each list *L* for genes that overlap with each gene set in the database^63,64^. The overlap or enrichment was assessed by weighted scoring based on mean correlation (r-value), and the enriched gene set received a normalized enrichment score, a significance *p*-value, and a FDR *q*-value. From the top positively and negatively correlated genes in each list *L*, we obtained separate sets *S*^+^ and *S*^−^ of positively and negatively enriched gene sets, respectively.

### Visualization of significantly enriched gene sets

We next visualized the key functional themes in these networks. For each pair of sets *S*^+^ and *S*^−^, we input gene set from GSEA into the Cytoscape network visualization software, and filtered for gene sets with FDR *q*-value < 0.05. We then used the Enrichment Map app to construct the gene set networks and annotated them with the Wordcloud extension^78–80^. This was done using the default settings except for a custom FDR *q*-value threshold of 0.05 (i.e., FDR < 0.05). This step returned four annotated enrichment maps for the list *L* of each cognitive function and for each of cortical and subcortical areas.

### Identifying candidate genes associated with memory

To identify candidate genes most likely to be relevant to the cognitive function, we identified genes frequently appearing across the gene sets with the leading edge analysis^63,64^ (LEA). For the analysis of each cognitive function and each ROI, we input the respective gene sets below FDR *q*-value < 0.05, after which the LEA identified the genes that appeared frequently across the leading-edge subset genes across gene sets in *S*^+^ or *S*^− 63,81^. The top-10 genes appearing most frequently in the positively and negatively enriched gene sets were designated as the candidate gene list *CL*. The outputs were eight candidate gene-cognition association lists *CL* of 10 genes each for all *S*^+^ and *S*^−^.

### Literature review of genetic signatures

We conducted a literature review and counted the number of ‘hits’ for the target cognitive function and the control function. To catalogue the function of each gene in cognition, we conducted a literature review on the candidate genes within each *CL* for its association with memory and motor function. This was done by reviewing experimental literature on Google Scholar, via a search query: (“gene name” AND (“memory” OR “amnesia” OR “Alzheimer’s”)), and (“gene name” AND (“motor” OR “ataxia” OR “Parkinson’s” OR “Huntington’s”)) respectively. The disorders were selected for keyword search because they related with deficiencies in memory and motor functioning. Causal evidence (in-vivo gene manipulations, mutants and pharmacological interventions) was classified as strong, while correlational evidence (computational gene associations, in vitro studies, differential gene expression studies and human case studies) was classified as weak. Literature evidence only counted as validation if it implicated the corresponding brain area, i.e., cortical or sub-cortical. Evidence of a given gene’s role solely in the non-analyzed brain region was not counted. For example, if a paper showed that the knockout of Gene A in the subcortex leads to disrupted memory, it would not count as evidence for the analysis of cortical memory.

### Assessing method performance in identifying candidate genes

We quantified method performance by significance testing and specificity assessment based on the prior literature review. We calculated the chance probability of obtaining *N* memory genes per gene list (without replacement) by calculating the proportion of known memory genes out of all 16906 genes (**Supplementary Table 3**). The same was done for motor function. The number of memory-related genes and motor-related genes were 614 and 78 respectively. These memory-related genes (memory or memory-related disorder) were compiled from three sources: (a) the literature review above, (b) the biological function gene sets 'GO:0007611 Learning or memory’, from database AmiGO2^82^ (version 2.4.26, release date 2016-08) and van Cauwenberghe *et al*. (2016)^83^. The motor-related genes (motor or motor-related disorder) were obtained from (a) the literature review above, (b) the biological function gene sets 'GO:0061743 Motor learning’ and 'GO:0061744 Motor behavior’ from database AmiGO2^82^ and (c) Lin *et al*. (2014)^84^. For example, if ten genes are selected randomly without replacement, the probability of all of them being memory-related genes is 3.72×10^−15^. With this, we calculated the probability of obtaining zero through ten memory-related genes (**Supplementary Table 3**).

### Dissociation genetic signatures of memory and motor function

The method specificity was determined by scoring the ratio of ‘hits’ versus ‘misses’. For example, in the memory analysis ‘hits’ correspond to gene-memory associations supported by literature, and ‘misses’ correspond to supported gene-motor function links. The larger the ratio of ‘hits’ over ‘misses’, the greater the method specificity. The evidence was weighted such that strong evidence ‘hits’ and weak evidence ‘hits’ received a full point and half-point, respectively. To quantify the method’s specificity, for each candidate gene list *CL*, we used the equations below for each memory and motor function *CL*s. If the method is specific, for memory analyses the memory specificity score should be above 50% and motor score below 50%, and vice versa.

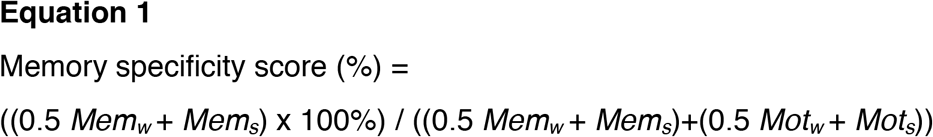

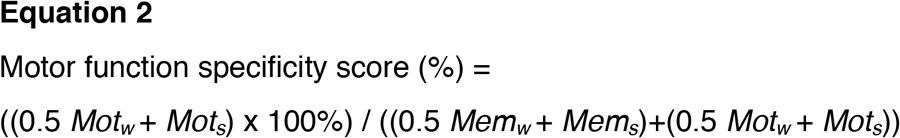

Legend:

*Mem_s_* = number of genes with strong evidence for its association with memory
*Mem_w_* = number of genes with weak evidence for its association with memory
*Mot_s_* = number of genes with strong evidence for its association with motor function
*Mot_w_* = number of genes with weak evidence for its association with motor function

## RESULTS

### Spatial correlation of transcriptome atlas and memory map

To analyze the spatial similarity between transcriptome and neuroimaging map, we used the Allen Human Brain Atlas (AHBA) and Neurosynth memory association map respectively. The AHBA was derived from six donor brains, and represented human brain gene expression in the left cortical and subcortical regions (N = 6) (Fig. 1a). The Neurosynth memory association map was a meta-study map (N = 2744) which represented of each brain region’s relevance for memory, specified by positive *z*-scores (Fig. 1a). We co-registered both maps into a common Montreal Neurological Institute 152 (MNI152) space. The areas in the memory association map were used to define the usable AHBA samples for the subsequent spatial correlation analysis. We conducted the spatial similarity analysis between AHBA and Neurosynth association map separately for cortical and subcortical regions, and for memory and motor function (for an example visualization, see **Fig. 2**). Genes with high spatial similarity between their expression map and memory term map are likely related to memory^15^.

**Figure 2.**
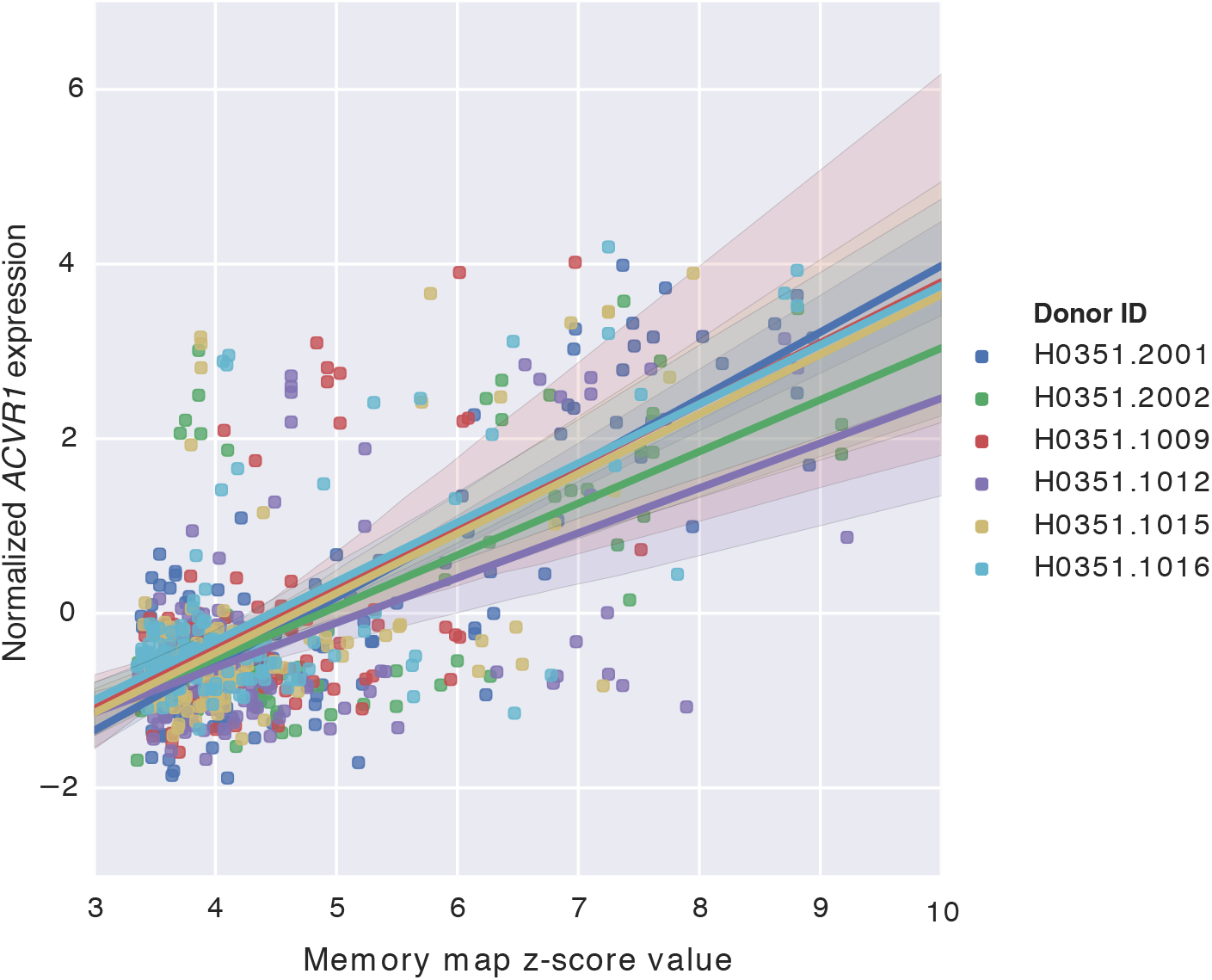
An example of spatial similarity analysis output. The expression levels for the top-correlated *ACVR1* gene are visualized as a function of the *z*-score of each voxel’s relevance to memory function. Normalized gene expression (*y*-axis) plotted against neuroimaging map *z*-scores (*x*-axis). Each colored regression line represents the best-fit line for each of six donors (colors); the translucent band around each line represents the 95% confidence interval estimate. (1.5-column image, colored)

Each analysis yielded a list *L*, which contained the mean correlation values of 16906 genes used for subsequent ranking (**Fig. 1b, Supplementary Table 1**). The mean correlation values were stable across donors (average *SD* = 0.122±0.03). The top-ten positively and negatively correlated genes for the memory cortical and subcortical analyses are shown in **Table 1**. There were more negatively correlated genes than positively correlated genes for both cortical and subcortical analyses of memory (**Supplementary Table 1**). We found 6770 positively and 10136 negatively correlated genes for the cortical areas, and positively 7256 and negatively 9649 genes for the subcortical areas. A positive correlation indicates higher gene expression in areas relevant for memory, and thus may be involved in enabling memory function^85^. A negative correlation implies the opposite, and the gene may play an inhibitory role^86^.

**Table 1.** Spatial similarity analysis output for memory function. Top-ten genes from each four lists *L* are ranked by the mean correlation values across six donor brains. The positively and negatively correlated genes are listed separately for cortical and subcortical areas.

| Correlation polarity | Cortical analysis |  | Subcortical analysis |  |
| --- | --- | --- | --- | --- |
|  | Gene | Mean *r* | Gene | Mean *r* |
| + | *ACVR1* | 0.64 | *MEF2C* | 0.43 |
|  | *TMTC1* | 0.61 | *YWHAH* | 0.42 |
|  | *SPHKAP* | 0.61 | *MAMLD1* | 0.41 |
|  | *SUSD4* | 0.61 | *KIAA1045* | 0.39 |
|  | *SLC1A1* | 0.59 | *CBX6* | 0.38 |
|  | *PPM1E* | 0.59 | *CLSPN* | 0.38 |
|  | *OVCH1* | 0.58 | *BSN* | 0.38 |
|  | *SMPD3* | 0.58 | *HMGA1* | 0.37 |
|  | *LINC00158* | 0.57 | *SYN2* | 0.36 |
|  | *MAMLD1* | 0.57 | *ABRACL* | 0.36 |
| – | *CITED2* | –0.60 | *HIST2H3D* | –0.35 |
|  | *TYRP1* | –0.60 | *LEMD1* | –0.36 |
|  | *MCHR2* | –0.61 | *PGM5* | –0.36 |
|  | *RORB* | –0.61 | *RPL11* | –0.36 |
|  | *TESPA1* | –0.61 | *C3orf58* | –0.36 |
|  | *RXFP1* | –0.62 | *HMGN2P46* | –0.37 |
|  | *KCNS1* | –0.62 | *DAO* | –0.38 |
|  | *CPNE9* | –0.62 | *H3F3C* | –0.38 |
|  | *SPARCL1* | –0.62 | *CRNDE* | –0.38 |
|  | *FSTL4* | –0.64 | *H3F3B* | –0.39 |

### Distinct biological processes of cortical-subcortical memory

To identify and characterize sets of genes that work towards a common biological function, we analyzed each of the cortical and subcortical lists *L* with GSEA-Preranked (Fig. 1c). This yielded positively scoring and negatively scoring gene sets, derived from the positively and negatively correlated genes of L, respectively. These gene sets were then grouped into functionally related clusters, and automatically annotated with biological themes. The complete GSEA results with the enriched biological processes and respective genes are available in **Supplementary Data 1**. Overall, the cortex and subcortex had distinct biological themes that were previously linked to memory. The cortex was linked to memory-related biological themes of associative learning, cellular respiration^87–89^, and protein targeting (**Fig. 3a**). The subcortex was linked memory-linked biological themes of the modulation of synaptic signaling^7^, neuronal recognition, adenylate cyclase coupling^90^, phosphatase activity^91–93^, limbic system development^94^, ribosome biogenesis^95^, mRNA transcription^96^, post-transcriptional processes^97,98^, epigenetics^99^, histone methylation^100,101^, and protein ubiquination^102,103^ (**Fig. 3b**).

**Figure 3.**
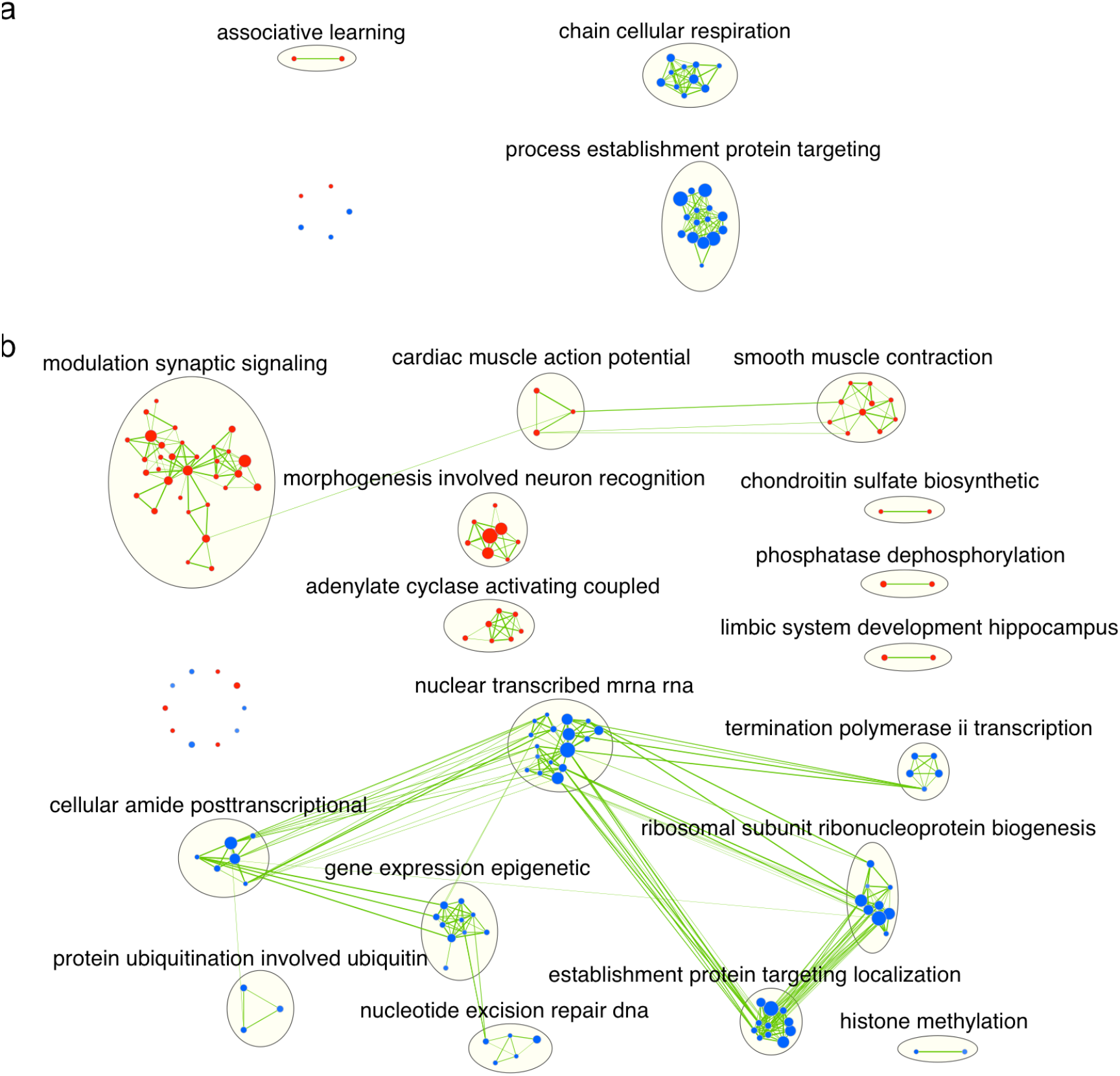
A functional map of memory. Significantly enriched gene sets (FDR q-value < 0.05) were visualized of with Enrichment Map in Cytoscape. Red nodes are positively scoring gene sets from *S*^+^ containing positively correlated genes, and vice versa. Each node is a gene set; the node size is proportional to the number of genes within the gene set. Edges represent shared genes between two nodes, edge-thickness represents the number of shared genes between nodes (minimum of 50%). Major functional groups of biologically related gene sets were clustered and circled. Clusters were automatically annotated with functional enrichment themes, which are the most frequently appearing terms within the cluster. (a) 35 gene sets associated with cortical areas for memory. (b) 147 gene sets associated with subcortical areas for memory. (2-column image, colored)

Examining the top gene sets within these biological themes, we found biological processes relevant for memory function in both cortical and subcortical analyses. In the positively scoring gene sets of the cortical analysis, we found the expected gene set “Associative learning” to be significantly enriched (FDR < .05). The most significant hit was the “regulation of clathrin-mediated endocytosis” gene set (**Table 2**). Clathrin-mediated endocytosis is relevant to memory, notably in AMPAR endocytosis in long-term potentiation based memory^104–106^. Dysregulated clathrin-mediated endocytosis processes has also been found in AD patients and animal models, and its inhibition can prevent memory impairment in an AD mouse model^107,108^. In the negatively scoring gene sets of the cortical analysis, the top two significant hits were the “Establishment of protein localization to endoplasmic reticulum” and “Nuclear transcribed mRNA catabolic process nonsense mediated decay” gene sets (**Table 2**). Nonsense-mediated decay of mRNA is a post-transcriptional quality control process which leads to the decay of mRNA^109–111^. An example target of this process is the *Arc* mRNA. *Arc* is a known long term potentiation-induced protein that is synthesized after learning and involved in memory consolidation^98,112^. Inhibition of the nonsense-mediated decay process increases synaptic strength, and the dysregulation of nonsense-mediated decay is associated with memory dysfunction^110,113^.

**Table 2.** Top significantly enriched biological process gene sets from the cortical and subcortica analyses of memory function. The positively and negatively scoring gene sets are ranked by their NES magnitudes respectively. The normalized enrichment score is indicative of a particular gene set’s degree of enrichment. Abbreviations: ‘NES’: Normalized Enrichment Score, ‘GO’: Gene Ontology; ‘mRNA’ messenger ribonucleic acid.

| Analysis | NES | GO biological process gene set |
| --- | --- | --- |
| Memory Cortical | 2.30 | Regulation of clathrin-mediated endocytosis |
|  | 2.22 | Visual behavior |
|  | 2.17 | Regulation of long term neuronal synaptic plasticity |
|  | -2.45 | Establishment of protein localization to endoplasmic reticulum |
|  | -2.33 | Nuclear transcribed mRNA catabolic process nonsense-mediated decay |
|  | -2.32 | Translational initiation |
| Memory Subcortical | 2.47 | Regulation of synaptic vesicle transport |
|  | 2.41 | Regulation of neurotransmitter secretion |
|  | 2.37 | Regulation of neurotransmitter transport |
|  | -2.73 | Nuclear transcribed mRNA catabolic process nonsense-mediated decay |
|  | -2.65 | Translational initiation |
|  | -2.64 | Establishment of protein localization to endoplasmic reticulum |

In the positively scoring gene sets of the subcortical analysis, we found the expected gene sets “Learning” and “Regulation of synaptic plasticity” to be significantly enriched (FDR < .05). The most significant hit was the “regulation of synaptic vesicle transport” gene set (**Table 2**). The regulation of vesicle exocytosis and vesicle cycling controls neurotransmitter release, which underlies synaptic transmission. Importantly, this regulation of vesicular transport is known to play a role in memory and working memory^114,115^. In the negatively scoring gene sets of the subcortical analysis, the top two significant hits were the “Nuclear transcribed mRNA catabolic process nonsense mediated decay” and “translational initiation” gene sets (**Table 2**). The relevance of nonsense mediated decay of mRNA has been discussed above. The suppression of translation initiation may be the state of mRNA translation at rest, which is alleviated only under conditions of learning^116^. Alternatively, the suppression of protein synthesis and ribosomal biogenesis may be relevant for memory function^117–119^.

### Distinct candidate genes of cortical-subcortical memory

To identify candidate genes that are most likely linked to human memory function, we identified genes that drive the enrichment score of multiple gene sets obtained above (Fig. 1d)^63,81,120^. This was done by applying the Leading Edge Analysis to the positively and negatively scoring gene sets above, followed by selecting the top-10 genes appearing most frequently across the leading edge subsets of the gene sets (**Supplementary Table 2**). These genes were then validated with animal model literature, which are classified as strong or weak evidence supporting the link between the gene and memory function (Fig. 1e). Strong evidence consists of causal gene manipulation or drug treatment studies, e.g., gene knockout leading to memory alteration. Weak evidence encompasses non-causal correlational or computational studies, e.g., gene upregulation that correlates with enhanced memory performance.

The candidate gene lists for cortical and subcortical memory were found to be largely distinct and are shown in **Table 3** and **4** (overlap = 6/40, **Supplementary Table 3**). For the positively correlated candidate genes of cortical memory areas, all ten had strong evidence supporting their role in memory function (**Table 3, Supplementary Table 3**). For example, *GRIN1* and *NTS* are causally linked to memory^121–123^. For the corresponding negatively correlated genes, nine out of ten had weak evidence for their role in memory (**Table 3**). These included ribosomal genes *RPS5* and *RPS13* which were differentially expressed in Alzheimer’s Disease models^124,125^, and *RPS16* and *RPS25* in long term memory^126,127^.

**Table 3.** Candidate gene lists from cortical analyses of memory. Candidate gene lists for the memory analysis of cortical regions, from positively and negatively correlated gene lists. Genes are ranked by the number of leading edge subsets they appear in, and subsequently by mean *r*-value. Abbreviations: ‘*CL*’: candidate gene list; ‘# leading edge subsets’: number of leading edge subsets that the gene was found in; ‘Mem_s_’: strong evidence for memory function; ‘Mem_w_’: weak evidence for memory function; ‘Mot_s_’: strong evidence for motor function; ‘Mot_w_’: weak evidence for motor function; ‘NA’: evidence for neither memory nor motor function; ‘+’: positively correlated candidate gene list; ‘−’: negatively correlated candidate gene list.

| CL | Gene | # leading edge subsets | mean *r* | Associated cognitive function |  |  |  |
| --- | --- | --- | --- | --- | --- | --- | --- |
|  |  |  |  | Mem<sub>s</sub> | Mem<sub>w</sub> | Mot<sub>s</sub> | Mot<sub>w</sub> |
| Memory Cortical + | *NPTN* | 3 | 0.48 | ✓<sup>142</sup> |  |  |  |
|  | *KIT* | 3 | 0.39 | ✓<sup>143</sup> |  |  |  |
|  | *SYNGAP1* | 3 | 0.36 | ✓<sup>144</sup> |  | ✓<sup>144</sup> |  |
|  | *NETO1* | 3 | 0.31 | ✓<sup>145</sup> |  |  |  |
|  | *GRIN1* | 3 | 0.25 | ✓<sup>121</sup> |  | ✓<sup>146</sup> |  |
|  | *NF1* | 3 | 0.23 | ✓<sup>147,148</sup> |  | ✓<sup>149</sup> |  |
|  | *NTS* | 2 | 0.51 | ✓<sup>122,123</sup> |  |  |  |
|  | *TANC1* | 2 | 0.46 | ✓<sup>150</sup> |  |  |  |
|  | *ATP1A3* | 2 | 0.45 | ✓<sup>151</sup> |  | ✓<sup>151</sup> |  |
|  | *NLGN3* | 2 | 0.41 | ✓<sup>152</sup> |  |  |  |
| Memory Cortical - | *RPL41* | 14 | -0.51 |  | ✓<sup>153,154</sup> |  |  |
|  | *RPL9* | 14 | -0.50 |  | ✓<sup>124,127</sup> |  |  |
|  | *RPS25* | 14 | -0.48 |  | ✓<sup>126</sup> |  |  |
|  | *RPS5* | 14 | -0.47 |  | ✓<sup>125</sup> |  | ✓<sup>155</sup> |
|  | *RPL37A* | 14 | -0.47 |  |  |  |  |
|  | *RPS16* | 14 | -0.46 |  | ✓<sup>127</sup> |  | ✓<sup>155</sup> |
|  | *RPL11* | 14 | -0.45 |  | ✓<sup>130</sup> |  | ✓<sup>156</sup> |
|  | *RPL18* | 14 | -0.45 |  | ✓<sup>157</sup> |  |  |
|  | *RPS13* | 14 | -0.44 |  | ✓<sup>125,158</sup> |  | ✓<sup>155</sup> |
|  | *RPS15* | 14 | -0.44 |  | ✓<sup>129,159</sup> |  | ✓<sup>160</sup> |

**Table 4.**
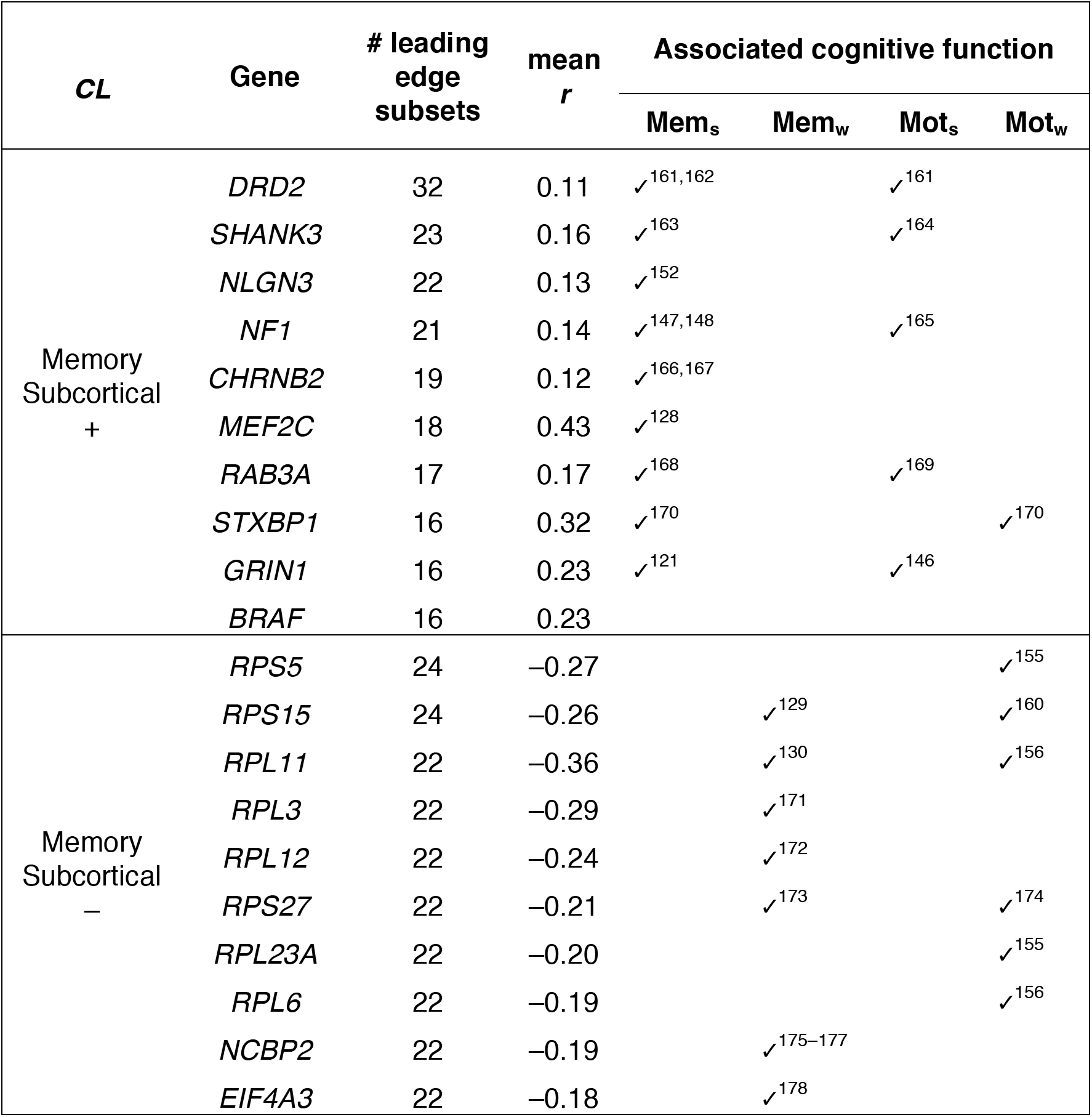
Candidate gene lists from subcortical analyses of memory. See Table 3 for notation.

For the positively correlated candidate genes of subcortical memory areas, nine out of ten had strong evidence supporting their role in memory function (**Table 4, Supplementary Table 3**). For example, *GRIN1* and *MEF2C* had causal links to memory^121,128^. For the corresponding negatively correlated candidate genes, seven out of ten had weaker evidence supporting their role in memory. These included *RPL11* and *RPS15* which were implicated in Alzheimer’s Disease^129,130^. Additionally, translation initiation factor *EIF4A3* and mRNA binding protein *NCBP2* were also implicated in striatal-learning and *d*-serine treatment associated with improved recognition and working memory respectively.

### Dissociable genetic signatures of memory and motor function

We assessed the effectiveness and specificity of the framework in identifying candidate genes for memory by using motor function as the control function (Fig. 1f). Generally, the framework was effective in identifying candidate genes for memory and motor function, inferred from the number of genes related to memory and motor function. For the memory analyses, we found that the number of genes associated with memory function were 7, 9, 9, and 10 for subcortical(−), subcortical(+), cortical(−), and cortical(+) candidate gene lists, respectively (**Supplementary Figure 1**). The probability of selecting seven memory-related genes by chance (i.e. from a catalogue of known memory-related genes) was small (*p* = 8.06×10^−11^) (**Supplementary Table 3 & 4**). For the motor analyses, the number of genes associated with motor function were 10, 9, 4 and 8 for subcortical(−), subcortical(+), cortical(−), and cortical(+) candidate gene lists respectively (**Supplementary Figure 1**). The probability of obtaining four motor function genes by chance was also low (*p* = 4.19×10^−10^) (**Supplementary Table 3 & 4**). The nine possible permutations of gene-cognition links from two cognitive functions and three strengths of evidence (strong, weak, none) are detailed and visualized in **Supplementary Table 3** and **Supplementary Figure 1** respectively. There was relatively little overlap between the candidate genes lists of memory and motor function (overlap=11/80, **Supplementary Table 3**). This overlap between memory and motor function gene lists may reflect the dual roles of the same gene in both memory and motor function, i.e. *RPL41* shared between memory cortical(−) and motor cortical(−) gene lists.

Generally, method specificity was high as more memory analysis genes were linked to memory function (i.e., high memory specificity score) as compared to motor function (i.e., low motor specificity score), and vice versa for motor function analysis. We found that the memory function *CLs* scored an average of 62.6% and 35.1% for the memory and motor function specificity scores, respectively (**Fig. 4, Table 5, Equations 1 and 2**). The motor function *CL*s scored 64.9% and 37.4% for the motor and memory function specificity scores, respectively. However, the *CL* containing negatively correlated genes from the motor function subcortical analysis scored 50% for both specificity scores (**Fig. 4, Table 5**), i.e., was non-specific. A multiple linear regression model confirmed that the cognitive function was the only statistically significant predictor of the specificity score (*t*(7) = −3.33, *p* < 0.05). These results suggest that the framework is relatively specific in identifying candidate genes associated with given cognitive functions.

**Figure 4.**
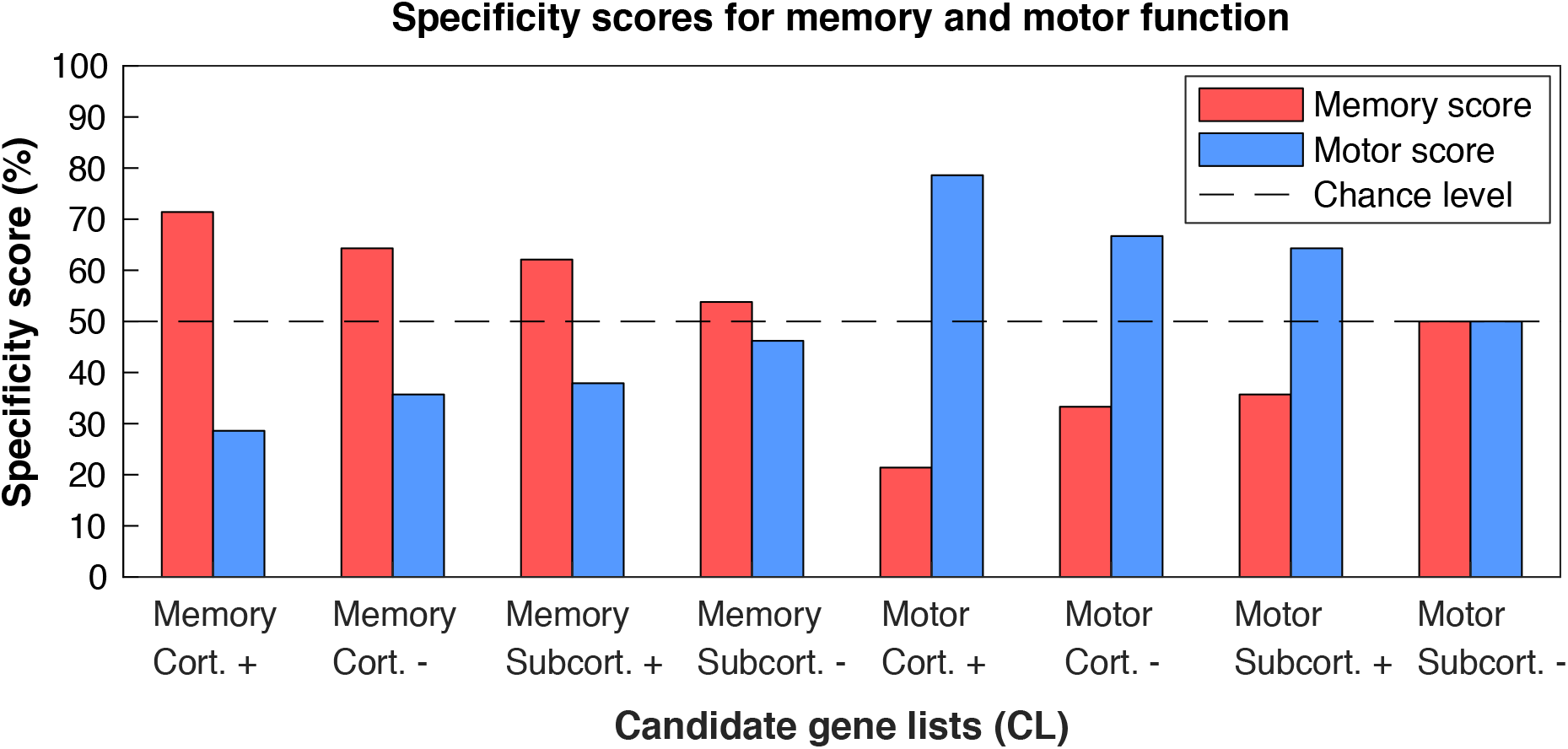
Performance assessment of candidate gene lists from cortical and subcortical analyses of memory and motor functions. The four memory gene lists are on the left, and vice versa for the motor function gene lists. The blue (memory) and red (motor) bars reflect the specificity score (*y*-axis) for each candidate gene list (*x*-axis). Horizontal solid line indicates the baseline method specificity score of 50%. The specificity of a given candidate gene list was higher than 50% for its corresponding function. Method specificity is scored with **Equations 1 and 2** (see methods). Abbreviations: ‘Cort.’: left cortical analysis, ‘Subcort.’: left subcortical analysis; ‘+’: positively correlated candidate gene list; ‘−’: negatively correlated candidate gene list. (2-column image, colored)

**Table 5.** Method specificity scores for candidate gene lists from cortical and subcortical analyses of memory and motor function. Candidate gene lists are scored by the method specificity equations 1 and 2 (see methods). If the method is specific, the specificity score for the target function should be >50%, and <50% for the control function; e.g. for memory analysis, memory is the target function and motor is the control function, and vice versa. Abbreviations: ‘*CL*’: candidate gene list; ‘+’: positively correlated candidate gene list; ‘−’: negatively correlated candidate gene list.

| Cognitive function | CL | Memory specificity score (%) | Motor function specificity score (%) |
| --- | --- | --- | --- |
| Memory | Cortical + | 71.4 | 28.6 |
|  | Cortical - | 64.3 | 35.7 |
|  | Subcortical + | 62.1 | 37.9 |
|  | Subcortical - | 53.8 | 46.2 |
| Motor function | Cortical + | 21.4 | 78.6 |
|  | Cortical - | 33.3 | 66.7 |
|  | Subcortical + | 35.7 | 64.3 |
|  | Subcortical - | 50.0 | 50.0 |

## DISCUSSION

Taken together, our results show that cortical and subcortical regions involved in human memory possess distinct genetic signatures. These genetic signatures are in agreement with prior research in animal models of memory, and were dissociable between memory and motor functions. This finding contributes to our knowledge of the functional differences of cortical and subcortical regions in healthy memory function and memory disorders. First, the distinct biological processes and candidate genes of cortical and subcortical memory regions may be related to the functional distinction in human cortical-subcortical memory. Second, animal memory literature supports the link between these genetic signatures and human memory in the cortex and subcortex. Third, the dissociation between memory and motor function candidate gene lists suggest that these genetic signatures are specific to each cognitive function. Thus, the strong similarities between the spatial patterns of human brain transcriptome and the functional neuroimaging map of memory can be exploited to identify molecular mechanisms of human memory and candidate biological processes/genes for drug development. Importantly, a drug discovery approach centered on human genomics^41,42^ may complement current efforts in identifying drug candidates for memory enhancement or the treatment of specific cortically- or subcortically-based memory disorders^44–46^.

Presently, most human memory evidence are derived from popular non-invasive methods such as Genome-Wide Association Studies^131^ (GWAS), which identifies links between gene variants and cognition^68^. GWAS was useful in discovery of a number of associations between genes and cognitive traits, such as *L1CAM* and repetition-based memory improvement^68,69,132,133^. However, GWAS ignores the spatially distributed gene expression in the brain by solely analyzing gene variants in relation to brain or behavioral measures^8,134^. Our framework overcomes these issues by accounting for spatial gene expression and identifying biological processes related to human memory. In line with this, we found that cortical and subcortical memory are associated with largely distinct memory-related processes, such as cortical memory and cellular respiration^87–89^, and subcortical memory and ribosome biogenesis^95^ (**Fig. 3a, 3b**). Furthermore, these findings could provide new insights into human memory. For example, we found epigenetic and histone methylation processes associated with human subcortical memory, which were previously linked to memory in rodents^100^. Modulating epigenetic regulators has previously shown promise in reversing Alzheimer’s disease in flies^135^.

Our framework also identifies candidate genes that drive the score of such memory-related biological processes, which are similarly distinct across cortical and subcortical memory. Within memory function, we found that the candidate genes of cortical and subcortical memory were largely distinct (**Table 3 and 4, Supplementary Table 3**). Interestingly, among positively correlated genes, *NLGN3, NF1* and *GRIN1* genes were common between cortical(+) and subcortical(+) gene lists. Among negatively correlated genes, *RPS5, RPL11* and *RPS15* overlapped between cortical(−) and subcortical(−) gene lists (**Table 3 and 4, Supplementary Table 3**). Overall, some genes were shared across cortical and subcortical memory brain regions with the same correlation polarity, and may play a general role in memory irrespective of brain region. Conversely, genes that are not shared across cortex and subcortex may play a more local role in memory function.

Across memory and motor function, candidate genes lists were generally distinct, indicating the relative specificity of the method. Shared genes consisted of ribosomal subunit protein genes, including *RPL41* shared between memory cortical(−) and motor cortical(−) gene lists. Intriguingly, multiple ribosomal subunit protein genes were shared between inversely correlated memory and motor function gene lists (**Table 3 and 4, Supplementary Table 3**). For example, *RPL11, RPL12, RPL23A, RPS5, RPS15* and *RPS27* were common between memory subcortical(−) and motor subcortical(+) gene lists. *RPL11, RPL12, RPS5* and *RPS15* overlapped between memory cortical(−) motor subcortical(+) gene lists (**Table 3 and 4, Supplementary Table 3**). The same ribosomal genes with inverse correlations implies differences in regulating ribosomal machinery depending on cognitive function. This suggests dual roles in enabling one function and inhibiting the other. This is in line with recent animal research, where recognition memory accompanies transcriptomic regulation^136^. In memory, it appears that ribosomal mRNA production is suppressed. While it may seem counterintuitive, as memory requires greatly upregulated protein production for memory formation^103,137,138^, the suppression of translational machinery mRNA may be the brain’s state at rest. This state may be reversed only under conditions of learning to enable increased memory protein production capacity^116^, i.e., increased ribosomal mRNA production and translation. Alternatively, this may also suggest that the suppression of protein synthesis and ribosomal biogenesis is relevant for memory function^117–119^. In the context of memory function, the identified biological processes and genes are related to memory in a convergent manner. These exist in both the cortical and subcortical regions, and recapitulate a trend of tight regulation of mRNA and synaptic plasticity in memory^136^. Researchers should thus view candidate genes in the context of biological processes to gain a comprehensive understanding of mechanisms underlying a given cognitive function.

This approach has downsides, as it is limited by the spatial resolution of both the cognitive function maps and human brain transcriptome. The sensitivity and statistical power of our framework will grow as the spatial resolution and sample size of the AHBA database increases. This is because the AHBA transcriptome map has a lower resolution compared to functional imaging map, especially in the cortex^71^. Furthermore, as the translation of gene mRNA into a functional product is heavily regulated, donor brain proteomes may be complementary in identifying genes linked to memory^139–141^.

## CONCLUSIONS

Here, using the Allen Institute brain transcriptional atlas and Neurosynth neuroimaging maps, we demonstrate that cortical and subcortical memory regions have distinct genetic signatures. These genetic signatures provide novel biological processes and molecular targets for understanding of human memory function in health and disease. Crucially, our framework may be applicable to other cognitive functions or neuroimaging modalities from the ones demonstrated here to identify and understand the genetic signatures of cognitive functions.

## COMPETING INTERESTS STATEMENT

The authors declare that they have no competing financial interests.

## AUTHOR CONTRIBUTIONS

P.K.T., E.A. and P.J.H. conceived the study. P.K. performed the analyses with technical support and guidance with data interpretation from E.A. and P.J.H. P.K.T. and E.A. wrote the manuscript. All authors reviewed the manuscript.

## ACKNOWLEDGEMENTS

Data were provided by the Allen Institute for Brain Science and the Neurosynth repository.

## FUNDING

This research did not receive any specific grant from funding agencies in the public, commercial, or not-for-profit sectors.

## REFERENCES

1. LaBar, K. S. & Cabeza, R. Cognitive neuroscience of emotional memory. Nat. Rev. Neurosci. (2006). doi:10.1038/nrn1825

2. D’Esposito, M. & Postle, B. R. The Cognitive Neuroscience of Working Memory. Annu. Rev. Psychol. (2015). doi:10.1146/annurev-psych-010814-015031

3. Squire, L. R. & Wixted, J. T. The Cognitive Neuroscience of Human Memory Since H.M. Annu. Rev. Neurosci. (2011). doi:10.1146/annurev-neuro-061010-113720

4. Papassotiropoulos, A. & de Quervain, D. J. F. Genetics of human episodic memory: Dealing with complexity. Trends in Cognitive Sciences (2011). doi:10.1016/j.tics.2011.07.005

5. Green, A. E. et al. Using genetic data in cognitive neuroscience: From growing pains to genuine insights. Nature Reviews Neuroscience (2008). doi:10.1038/nrn2461

6. Atzmon, G., Bayer, A., Freudenberg-Hua, Y., Li, W. & Davies, P. The Role of Genetics in Advancing Precision Medicine for Alzheimer’s Disease—A Narrative Review. Front. Med. (2018). doi:10.3389/fmed.2018.00108

7. Kandel, E. R., Dudai, Y. & Mayford, M. R. The molecular and systems biology of memory. Cell (2014). doi:10.1016/j.cell.2014.03.001

8. Hawrylycz, M. J. et al. An anatomically comprehensive atlas of the adult human brain transcriptome. Nature 489, 391–399 (2012).

9. Yarkoni, T., Poldrack, R. A., Nichols, T. E., Van Essen, D. C. & Wager, T. D. Large-scale automated synthesis of human functional neuroimaging data. Nat. Methods 8, 665–70 (2011).

10. Romero-Garcia, R. et al. Structural covariance networks are coupled to expression of genes enriched in supragranular layers of the human cortex. Neuroimage (2018). doi:10.1016/j.neuroimage.2017.12.060

11. Ritchie, J., Pantazatos, S. P. & French, L. Transcriptomic characterization of MRI contrast with focus on the T1-w/T2-w ratio in the cerebral cortex. Neuroimage (2018). doi:10.1016/j.neuroimage.2018.03.027

12. Wang, G.-Z. et al. Correspondence between Resting-State Activity and Brain Gene Expression. Neuron 88, 659–66 (2015).

13. Krienen, F. M., Yeo, B. T. T., Ge, T., Buckner, R. L. & Sherwood, C. C. Transcriptional profiles of supragranular-enriched genes associate with corticocortical network architecture in the human brain. Proc. Natl. Acad. Sci. 113, E469–E478 (2016).

14. Vértes, P. E. et al. Gene transcription profiles associated with inter-modular hubs and connection distance in human functional magnetic resonance imaging networks. Philos. Trans. R. Soc. B Biol. Sci. (2016). doi:10.1098/rstb.2015.0362

15. Fox, A. S., Chang, L. J., Gorgolewski, K. J. & Yarkoni, T. Bridging Cognition and Genetics Using Large-scale Spatial Analysis of Neuroimaging and Neurogenetic Data. Biol. Psychiatry 77, 377S–377S (2015).

16. Diez, I. & Sepulcre, J. Neurogenetic profiles delineate large-scale connectivity dynamics of the human brain. Nat. Commun. 9, 3876 (2018).

17. Anderson, K. M. et al. Gene expression links functional networks across cortex and striatum. Nat. Commun. (2018). doi:10.1038/s41467-018-03811-x

18. Bonelli, R. M. & Cummings, J. L. Frontal-subcortical dementias. Neurologist (2008). doi:10.1097/NRL.0b013e31815b0de2

19. Cummings, J. L. Vascular subcortical dementias: Clinical aspects. Dement. Geriatr. Cogn. Disord. (1994). doi:10.1159/000106718

20. Schott, B. H. et al. Redefining implicit and explicit memory: The functional neuroanatomy of priming, remembering, and control of retrieval. Proc. Natl. Acad. Sci. (2005). doi:10.1073/pnas.0409070102

21. Squire, L. R. & Dede, A. J. O. Conscious and unconscious memory systems. Cold Spring Harb. Perspect. Biol. (2015). doi:10.1101/cshperspect.a021667

22. Stern, S. A. & Alberini, C. M. Mechanisms of memory enhancement. Wiley Interdiscip. Rev. Syst. Biol. Med. 5, 37–53 (2013).

23. Matthews, P. M. & Hampshire, A. Clinical Concepts Emerging from fMRI Functional Connectomics. Neuron 91, 511–528 (2016).

24. Davies, P. & Koppel, J. Mechanism-based treatments for Alzheimer’s disease. Dialogues Clin. Neurosci. 11, 159–69 (2009).

25. Solari, N., Bonito-Oliva, A., Fisone, G. & Brambilla, R. Understanding cognitive deficits in Parkinson’s disease: lessons from preclinical animal models. Learn. Mem. 20, 592–600 (2013).

26. Graf, P. & Schacter, D. L. Implicit and Explicit Memory for New Associations in Normal and Amnesic Subjects. J. Exp. Psychol. Learn. Mem. Cogn. (1985). doi:10.1037/0278-7393.11.3.501

27. Gaillard, R. et al. Converging intracranial markers of conscious access. PLoS Biol. (2009). doi:10.1371/journal.pbio.1000061

28. Squire, L. R., Genzel, L., Wisted, J. T. & Morris, R. G. M. Memory Consolidation. Cold Spring Harb. Perspect. Biol. (2015). doi:10.1037/a0037983.Hearing

29. Schroeter, M. L., Stein, T., Maslowski, N. & Neumann, J. Neural correlates of Alzheimer’s disease and mild cognitive impairment: A systematic and quantitative meta-analysis involving 1351 patients. Neuroimage (2009). doi:10.1016/j.neuroimage.2009.05.037

30. Guimond, S., Chakravarty, M. M., Bergeron-Gagnon, L., Patel, R. & Lepage, M. Verbal memory impairments in schizophrenia associated with cortical thinning. NeuroImage Clin. (2016). doi:10.1016/j.nicl.2015.12.010

31. Heckers, S. et al. Impaired recruitment of the hippocampus during conscious recollection in schizophrenia. Nat. Neurosci. (1998). doi:10.1038/1137

32. Molina, V., Taboada, D., Aragüés, M., Hernández, J. A. & Sanz-Fuentenebro, J. Greater clinical and cognitive improvement with clozapine and risperidone associated with a thinner cortex at baseline in first-episode schizophrenia. Schizophr. Res. (2014). doi:10.1016/j.schres.2014.06.042

33. Shaik, S. S. & Varma, A. R. Differentiating the dementias: A neurological approach. Progress in Neurology and Psychiatry (2012). doi:10.1002/pnp.224

34. Alloway, T. P., Rajendran, G. & Archibald, L. M. D. Working Memory in Children With Developmental Disorders. J. Learn. Disabil. (2009). doi:10.1177/0022219409335214

35. Park, D. C. & Festini, S. B. Theories of memory and aging: A look at the past and a glimpse of the future. Journals of Gerontology - Series B Psychological Sciences and Social Sciences (2017). doi:10.1093/geronb/gbw066

36. Kirova, A.-M., Bays, R. B. & Lagalwar, S. Working Memory and Executive Function Decline across Normal Aging, Mild Cognitive Impairment, and Alzheimer’s Disease. Biomed Res. Int. (2015). doi:10.1155/2015/748212

37. Ward, E. V, Berry, C. J. & Shanks, D. R. Age effects on explicit and implicit memory. Front Psychol (2013). doi:10.3389/fpsyg.2013.00639

38. Pittenger, C. Disorders of memory and plasticity in psychiatric disease. Dialogues Clin. Neurosci. (2013). doi:10.1038/jid.2015.269

39. Armario, A. & Nadal, R. Individual differences and the characterization of animal models of psychopathology: A strong challenge and a good opportunity. Frontiers in Pharmacology (2013). doi:10.3389/fphar.2013.00137

40. Keeler, J. F. & Robbins, T. W. Translating cognition from animals to humans. Biochemical Pharmacology (2011). doi:10.1016/j.bcp.2010.12.028

41. Papassotiropoulos, A. & de Quervain, D. J. F. Failed drug discovery in psychiatry: Time for human genome-guided solutions. Trends in Cognitive Sciences (2015). doi:10.1016/j.tics.2015.02.002

42. Breen, G. et al. Translating genome-wide association findings into new therapeutics for psychiatry. Nature Neuroscience (2016). doi:10.1038/nn.4411

43. Berto, S., Wang, G.-Z., Germi, J., Lega, B. C. & Konopka, G. Human Genomic Signatures of Brain Oscillations During Memory Encoding. Cereb. Cortex (2017). doi:10.1093/cercor/bhx083

44. Robbins, T. W. Cross-species studies of cognition relevant to drug discovery: a translational approach. British Journal of Pharmacology (2017). doi:10.1111/bph.13826

45. Wallace, T. L., Ballard, T. M. & Glavis-Bloom, C. Animal paradigms to assess cognition with translation to humans. Handb. Exp. Pharmacol. (2015). doi:10.1007/978-3-319-16522-6_2

46. Denayer, T., Stöhr, T. & Van Roy, M. Animal models in translational medicine: Validation and prediction. New Horizons Transl. Med. (2014). doi:10.1016/j.nhtm.2014.08.001

47. Zapala, M. a et al. Adult mouse brain gene expression patterns bear an embryologic imprint. Proc. Natl. Acad. Sci. U. S. A. (2005). doi:10.1073/pnas.0503357102

48. Grange, P. et al. Cell-type-based model explaining coexpression patterns of genes in the brain. Proc. Natl. Acad. Sci. (2014). doi:10.1073/pnas.1312098111

49. Kaufman, A., Dror, G., Meilijson, I. & Ruppin, E. Gene expression of Caenorhabditis elegans neurons carries information on their synaptic connectivity. PLoS Comput. Biol. (2006). doi:10.1371/journal.pcbi.0020167

50. French, L. & Pavlidis, P. Relationships between gene expression and brain wiring in the adult rodent brain. PLoS Comput. Biol. (2011). doi:10.1371/journal.pcbi.1001049

51. Bagot, R. C. C. et al. Circuit-wide Transcriptional Profiling Reveals Brain Region-Specific Gene Networks Regulating Depression Susceptibility. Neuron (2016). doi:10.1016/j.neuron.2016.04.015

52. Richiardi, J. et al. Correlated gene expression supports synchronous activity in brain networks. Science (80-.). 348, 1241–1244 (2015).

53. Cioli, C., Abdi, H., Beaton, D., Burnod, Y. & Mesmoudi, S. Differences in human cortical gene expression match the temporal properties of large-scale functional networks. PLoS One (2014). doi:10.1371/journal.pone.0115913

54. Whitaker, K. J. et al. Adolescence is associated with genomically patterned consolidation of the hubs of the human brain connectome. Proc. Natl. Acad. Sci. (2016). doi:10.1073/pnas.1601745113

55. Jung, K. et al. Effective connectivity during working memory and resting states: A DCM study. Neuroimage (2018). doi:10.1016/j.neuroimage.2017.12.067

56. Manelis, A. & Reder, L. M. Effective connectivity among the working memory regions during preparation for and during performance of the n-back task. Front. Hum. Neurosci. (2014). doi:10.3389/fnhum.2014.00593

57. Grady, C. L., McIntosh, A. R. & Craik, F. I. M. Age-related differences in the functional connectivity of the hippocampus during memory encoding. Hippocampus (2003). doi:10.1002/hipo.10114

58. Hampstead, B. M., Khoshnoodi, M., Yan, W., Deshpande, G. & Sathian, K. Patterns of effective connectivity during memory encoding and retrieval differ between patients with mild cognitive impairment and healthy older adults. Neuroimage (2016). doi:10.1016/j.neuroimage.2015.10.002

59. Ranganath, C., Heller, A., Cohen, M. X., Brozinsky, C. J. & Rissman, J. Functional connectivity with the hippocampus during successful memory formation. Hippocampus (2005). doi:10.1002/hipo.20141

60. Jeong, W., Chung, C. K. & Kim, J. S. Episodic memory in aspects of large-scale brain networks. Front. Hum. Neurosci. (2015). doi:10.3389/fnhum.2015.00454

61. Fjell, a. M. et al. Brain Events Underlying Episodic Memory Changes in Aging: A Longitudinal Investigation of Structural and Functional Connectivity. Cereb. Cortex (2015). doi:10.1093/cercor/bhv102

62. Ngo, C. T. et al. White matter structural connectivity and episodic memory in early childhood. Dev. Cogn. Neurosci. (2017). doi:10.1016/j.dcn.2017.11.001

63. Subramanian, A. et al. Gene set enrichment analysis: A knowledge-based approach for interpreting genome-wide expression profiles. Proc. Natl. Acad. Sci. 102, 15545–15550 (2005).

64. Mootha, V. K. et al. PGC-1?-responsive genes involved in oxidative phosphorylation are coordinately downregulated in human diabetes. Nat. Genet. 34, 267–273 (2003).

65. Thomassen, M., Tan, Q. & Kruse, T. A. Gene expression meta-analysis identifies metastatic pathways and transcription factors in breast cancer. BMC Cancer 8, 394 (2008).

66. Ersland, K. M. et al. Gene-Based Analysis of Regionally Enriched Cortical Genes in GWAS Data Sets of Cognitive Traits and Psychiatric Disorders. PLoS One 7, e31687 (2012).

67. Lee, M.-Y. et al. Alteration of Synaptic Activity–Regulating Genes Underlying Functional Improvement by Long-term Exposure to an Enriched Environment in the Adult Brain. Neurorehabil. Neural Repair 27, 561–574 (2013).

68. Heck, A. et al. Converging genetic and functional brain imaging evidence links neuronal excitability to working memory, psychiatric disease, and brain activity. Neuron 81, 1203–13 (2014).

69. Luksys, G. et al. Computational dissection of human episodic memory reveals mental process-specific genetic profiles. Proc. Natl. Acad. Sci. (2015). doi:10.1073/pnas.1500860112

70. Rizzo, G. et al. MENGA: A New Comprehensive Tool for the Integration of Neuroimaging Data and the Allen Human Brain Transcriptome Atlas. PLoS One 11, e0148744 (2016).

71. Hawrylycz, M. et al. Multi-scale correlation structure of gene expression in the brain. Neural Networks 24, 933–942 (2011).

72. Volkow, N. D. et al. PET evaluation of the dopamine system of the human brain. J. Nucl. Med. 37, 1242–56 (1996).

73. Mengod, G. et al. Visualization of dopamine D1, D2 and D3 receptor mRNA’s in human and rat brain. Neurochem. Int. 20, 33–43 (1992).

74. Pappata, S. et al. In vivo detection of striatal dopamine release during reward: a PET study with [(11)C]raclopride and a single dynamic scan approach. Neuroimage 16, 1015–27 (2002).

75. Schott, B. H. et al. Mesolimbic Functional Magnetic Resonance Imaging Activations during Reward Anticipation Correlate with Reward-Related Ventral Striatal Dopamine Release. J. Neurosci. 28, 14311–14319 (2008).

76. D’Ardenne, K., McClure, S. M., Nystrom, L. E. & Cohen, J. D. BOLD Responses Reflecting Dopaminergic Signals in the Human Ventral Tegmental Area. Science (80-.). 319, 1264–1267 (2008).

77. Reich, M. et al. GenePattern 2.0. Nat. Genet. 38, 500–501 (2006).

78. Merico, D., Isserlin, R., Stueker, O., Emili, A. & Bader, G. D. Enrichment Map: A Network-Based Method for Gene-Set Enrichment Visualization and Interpretation. PLoS One 5, e13984 (2010).

79. Cline, M. S. et al. Integration of biological networks and gene expression data using Cytoscape. Nat. Protoc. 2, 2366–2382 (2007).

80. Oesper, L., Merico, D., Isserlin, R. & Bader, G. D. WordCloud: a Cytoscape plugin to create a visual semantic summary of networks. Source Code Biol. Med. 6, 7 (2011).

81. Fleming, D. S. & Miller, L. C. Leading edge analysis of transcriptomic changes during pseudorabies virus infection. Genomics data 10, 104–106 (2016).

82. Carbon, S. et al. AmiGO: online access to ontology and annotation data. Bioinformatics 25, 288–9 (2009).

83. Van Cauwenberghe, C., Van Broeckhoven, C. & Sleegers, K. The genetic landscape of Alzheimer disease: clinical implications and perspectives. Genet. Med. 18, 421–430 (2016).

84. Lin, M. K. et al. Genetics and genomics of Parkinson’s disease. Genome Med. 6, 48 (2014).

85. Wang, D. et al. Genetic enhancement of memory and long-term potentiation but not CA1 long-term depression in NR2B transgenic rats. PLoS One (2009). doi:10.1371/journal.pone.0007486

86. Gai, Y., Liu, Z., Cervantes-Sandoval, I. & Davis, R. L. Drosophila SLC22A Transporter Is a Memory Suppressor Gene that Influences Cholinergic Neurotransmission to the Mushroom Bodies. Neuron (2016). doi:10.1016/j.neuron.2016.03.017

87. Rojas, J. C., Bruchey, A. K. & Gonzalez-Lima, F. Neurometabolic mechanisms for memory enhancement and neuroprotection of methylene blue. Progress in Neurobiology (2012). doi:10.1016/j.pneurobio.2011.10.007

88. Riha, P. D., Bruchey, A. K., Echevarria, D. J. & Gonzalez-Lima, F. Memory facilitation by methylene blue: Dose-dependent effect on behavior and brain oxygen consumption. Eur. J. Pharmacol. (2005). doi:10.1016/j.ejphar.2005.02.001

89. Grimm, A. & Eckert, A. Brain aging and neurodegeneration: from a mitochondrial point of view. Journal of Neurochemistry (2017). doi:10.1111/jnc.14037

90. Livingstone, M. S., Sziber, P. P. & Quinn, W. G. Loss of calcium/calmodulin responsiveness in adenylate cyclase of rutabaga, a Drosophila learning mutant. Cell (1984). doi:10.1016/0092-8674(84)90316-7

91. Braithwaite, S. P., Stock, J. B., Lombroso, P. & Nairn, A. Protein Phosphatases and Alzheimers’s Disease. Prog Mol Biol Trasnsl Sci (2012). doi:10.1016/B978-0-12-396456-4.00012-2.Protein

92. Sweatt, J. D. Memory mechanisms: The yin and yang of protein phosphorylation. Curr. Biol. (2001). doi:10.1016/S0960-9822(01)00216-0

93. Woolfrey, K. M. & Dell’Acqua, M. L. Coordination of protein phosphorylation and dephosphorylation in synaptic plasticity. J. Biol. Chem. (2015). doi:10.1074/jbc.R115.657262

94. Deal, A. L., Erickson, K. J., Shiers, S. I. & Burman, M. A. Limbic system development underlies the emergence of classical fear conditioning during the third and fourth weeks of life in the rat. Behav. Neurosci. (2016). doi:10.1037/bne0000130

95. Hernández, A. I., Alarcon, J. M. & Allen, K. D. New ribosomes for new memories? Commun. Integr. Biol. 8, e1017163 (2015).

96. Alberini, C. M. & Kandel, E. R. The regulation of transcription in memory consolidation. Cold Spring Harb. Perspect. Biol. (2015). doi:10.1101/cshperspect.a021741

97. Rosi, S. Neuroinflammation and the plasticity-related immediate-early gene Arc. Brain, Behavior, and Immunity (2011). doi:10.1016/j.bbi.2011.02.003

98. Bramham, C. R. et al. The Arc of synaptic memory. Experimental Brain Research 200, 125–140 (2010).

99. Zovkic, I. B., Guzman-Karlsson, M. C. & Sweatt, J. D. Epigenetic regulation of memory formation and maintenance. Learn. Mem. 20, 61–74 (2013).

100. Kim, S. & Kaang, B.-K. Epigenetic regulation and chromatin remodeling in learning and memory. Exp. Mol. Med. (2017). doi:10.1038/emm.2016.140

101. Gupta, S. et al. Histone Methylation Regulates Memory Formation. J. Neurosci. (2010). doi:10.1523/JNEUROSCI.3732-09.2010

102. Jarome, T. J. & Devulapalli, R. K. The Ubiquitin-Proteasome System and Memory: Moving Beyond Protein Degradation. Neuroscientist (2018). doi:10.1177/1073858418762317

103. Jarome, T. J. & Helmstetter, F. J. Protein degradation and protein synthesis in long-term memory formation. Front. Mol. Neurosci. 7, 61 (2014).

104. Man, Y. H. et al. Regulation of AMPA receptor-mediated synaptic transmission by clathrin- dependent receptor internalization. Neuron 25, 649–662 (2000).

105. Lee, S. H. et al. Clathrin Adaptor AP2 and NSF Interact with Overlapping Sites of GluR2 and Play Distinct Roles in AMPA Receptor Trafficking and Hippocampal LTD. Neuron 36, 661–674 (1994).

106. Ahmadian, G. et al. Tyrosine phosphorylation of GluR2 is required for insulin-stimulated AMPA receptor endocytosis and LTD. EMBO J. 23, 1040–1050 (2004).

107. Cao, Y., Xiao, Y., Ravid, R. & Guan, Z. Z. Changed clathrin regulatory proteins in the brains of Alzheimer’s disease patients and animal models. J. Alzheimer’s Dis. 22, 329–342 (2010).

108. Kuboyama, T., Lee, Y. A., Nishiko, H. & Tohda, C. Inhibition of clathrin-mediated endocytosis prevents amyloid beta-induced axonal damage. Neurobiol Aging 36, 1808–1819 (2015).

109. Amrani, N., Sachs, M. S. & Jacobson, A. Early nonsense: mRNA decay solves a translational problem. Nature Reviews Molecular Cell Biology 7, 415–425 (2006).

110. Giorgi, C. et al. The EJC Factor eIF4AIII Modulates Synaptic Strength and Neuronal Protein Expression. Cell 130, 179–191 (2007).

111. Peebles, C. L. & Finkbeiner, S. RNA decay back in play. Nature Neuroscience 10, 1083–1084 (2007).

112. Steward, O., Wallace, C. S., Lyford, G. L. & Worley, P. F. Synaptic activation causes the mRNA for the leg Arc to localize selectively near activated postsynaptic sites on dendrites. Neuron 21, 741–751 (1998).

113. Nguyen, L. S. et al. Contribution of copy number variants involving nonsense-mediated mRNA decay pathway genes to neuro-developmental disorders. Hum. Mol. Genet. 22, 1816–25 (2013).

114. Cottrell, J. R. et al. Working memory impairment in calcineurin knock-out mice is associated with alterations in synaptic vesicle cycling and disruption of high-frequency synaptic and network activity in prefrontal cortex. J. Neurosci. 33, 10938–49 (2013).

115. Lin, R. C. & Scheller, R. H. Mechanisms of Synaptic Vesicle Exocytosis. Annu. Rev. Cell Dev. Biol. 16, 19–49 (2000).

116. Chesnokova, E., Bal, N. & Kolosov, P. Kinases of eIF2a Switch Translation of mRNA Subset during Neuronal Plasticity. Int. J. Mol. Sci. 18, (2017).

117. Capitano, F. et al. RNA polymerase I transcription is modulated by spatial learning in different brain regions. J. Neurochem. 136, 706–716 (2016).

118. Cho, J. et al. Multiple repressive mechanisms in the hippocampus during memory formation. Science 350, 82–7 (2015).

119. Levy, R., Levitan, D. & Susswein, A. J. New learning while consolidating memory during sleep is actively blocked by a protein synthesis dependent process. Elife 5, (2016).

120. Darby, M. M., Yolken, R. H. & Sabunciyan, S. Consistently altered expression of gene sets in postmortem brains of individuals with major psychiatric disorders. Transl. Psychiatry 6, e890 (2016).

121. Umemori, J. et al. ENU-mutagenesis mice with a non-synonymous mutation in Grin1 exhibit abnormal anxiety-like behaviors, impaired fear memory, and decreased acoustic startle response. BMC Res. Notes 6, 203 (2013).

122. Yamada, D., Wada, E., Amano, T., Wada, K. & Sekiguchi, M. Lack of neurotensin type 1 receptor facilitates contextual fear memory depending on the memory strength. Pharmacol. Biochem. Behav. 96, 363–369 (2010).

123. Xiao, Z. et al. Activation of Neurotensin Receptor 1 Facilitates Neuronal Excitability and Spatial Learning and Memory in the Entorhinal Cortex: Beneficial Actions in an Alzheimer’s Disease Model. J. Neurosci. 34, 7027–7042 (2014).

124. Kong, W. et al. Independent component analysis of Alzheimer’s DNA microarray gene expression data. Mol. Neurodegener. 4, 5 (2009).

125. Hernández-Ortega, K., Garcia-Esparcia, P., Gil, L., Lucas, J. J. & Ferrer, I. Altered Machinery of Protein Synthesis in Alzheimer’s: From the Nucleolus to the Ribosome. Brain Pathol. 26, 593–605 (2016).

126. Winbush, A. et al. Identification of Gene Expression Changes Associated With Long-Term Memory of Courtship Rejection in Drosophila Males. G3&58; Genes|Genomes|Genetics 2, 1437–1445 (2012).

127. Shea, C. J. A. et al. Variable impact of chronic stress on spatial learning and memory in BXD mice. Physiol. Behav. 150, 69–77 (2015).

128. Barbosa, A. C. et al. MEF2C, a transcription factor that facilitates learning and memory by negative regulation of synapse numbers and function. Proc. Natl. Acad. Sci. U. S. A. 105, 9391 (2008).

129. Wang, J. et al. Chromosome 19p in Alzheimer’s Disease: When Genome Meets Transcriptome. J. Alzheimer’s Dis. 38, 245–250 (2014).

130. Oka, S. et al. Human mitochondrial transcriptional factor A breaks the mitochondria-mediated vicious cycle in Alzheimer’s disease. Sci. Rep. 6, 37889 (2016).

131. Burton, P. R. et al. Genome-wide association study of 14,000 cases of seven common diseases and 3,000 shared controls. Nature 447, 661–678 (2007).

132. Milnik, A. et al. Association of KIBRA with episodic and working memory: A meta-analysis. Am. J. Med. Genet. Part B Neuropsychiatr. Genet. (2012). doi:10.1002/ajmg.b.32101

133. Papassotiropoulos, A. et al. A genome-wide survey and functional brain imaging study identify CTNNBL1 as a memory-related gene. Mol. Psychiatry (2013). doi:10.1038/mp.2011.148

134. Mahfouz, A., Huisman, S. M. H., Lelieveldt, B. P. F. & Reinders, M. J. T. Brain transcriptome atlases: a computational perspective. Brain Struct. Funct. 222, 1557–1580 (2017).

135. Panikker, P. et al. Restoring Tip60 HAT/HDAC2 balance in the neurodegenerative brain relieves epigenetic transcriptional repression and reinstates cognition. J. Neurosci. (2018). doi:10.1523/JNEUROSCI.2840-17.2018

136. Scott, H. et al. Recognition memory-induced gene expression in the perirhinal cortex: A transcriptomic analysis. Behav. Brain Res. (2017). doi:10.1016/j.bbr.2017.04.007

137. Hernandez, P. J. & Abel, T. The role of protein synthesis in memory consolidation: Progress amid decades of debate. Neurobiol. Learn. Mem. 89, 293–311 (2008).

138. Sacktor, T. C. How does PKMζ maintain long-term memory? Nat. Rev. Neurosci. 12, 9–15 (2011).

139. Park, Y. M. et al. Profiling human brain proteome by multi-dimensional separations coupled with MS. Proteomics 6, 4978–4986 (2006).

140. Sjöstedt, E. et al. Defining the Human Brain Proteome Using Transcriptomics and Antibody-Based Profiling with a Focus on the Cerebral Cortex. PLoS One 10, e0130028 (2015).

141. Lubec, G., Krapfenbauer, K. & Fountoulakis, M. Proteomics in brain research: potentials and limitations. Prog. Neurobiol. 69, 193–211 (2003).

142. Bhattacharya, S. et al. Genetically Induced Retrograde Amnesia of Associative Memories After Neuroplastin Ablation. Biol. Psychiatry 81, 124–135 (2017).

143. Katafuchi, T., Li, A. J., Hirota, S., Kitamura, Y. & Hori, T. Impairment of spatial learning and hippocampal synaptic potentiation in c-kit mutant rats. Learn. Mem. 7, 383–92

144. Muhia, M. et al. Molecular and behavioral changes associated with adult hippocampus-specific SynGAP1 knockout. Learn. Mem. 19, 268–281 (2012).

145. Ng, D. et al. Neto1 Is a Novel CUB-Domain NMDA Receptor–Interacting Protein Required for Synaptic Plasticity and Learning. PLoS Biol. 7, e1000041 (2009).

146. Kiefer, F. et al. Involvement of NMDA receptors in alcohol-mediated behavior: mice with reduced affinity of the NMDA R1 glycine binding site display an attenuated sensitivity to ethanol. Biol. Psychiatry 53, 345–51 (2003).

147. Silva, A. J. et al. A mouse model for the learning and memory deficits associated with neurofibromatosis type I. Nat. Genet. 15, 281–284 (1997).

148. Costa, R. M. et al. Learning deficits, but normal development and tumor predisposition, in mice lacking exon 23a of Nf1. Nat. Genet. 27, 399–405 (2001).

149. Feldmann, R., Denecke, J., Grenzebach, M., Schuierer, G. & Weglage, J. Neurofibromatosis type 1. Motor and cognitive function and T2-weighted MRI hyperintensities. Neurology 61, 1725–1728 (2003).

150. Han, S. et al. Regulation of Dendritic Spines, Spatial Memory, and Embryonic Development by the TANC Family of PSD-95-Interacting Proteins. J. Neurosci. 30, 15102–15112 (2010).

151. Holm, T. H. et al. Cognitive deficits caused by a disease-mutation in the α3 Na(+)/K(+)-ATPase isoform. Sci. Rep. 6, 31972 (2016).

152. Tabuchi, K. et al. A neuroligin-3 mutation implicated in autism increases inhibitory synaptic transmission in mice. Science 318, 71–6 (2007).

153. Wang, A. et al. Ribosomal protein RPL41 induces rapid degradation of ATF4, a transcription factor critical for tumour cell survival in stress. J. Pathol. 225, 285–292 (2011).

154. Fayaz, S. M. & Rajanikant, G. K. ATF4: the perpetrator in axonal-mediated neurodegeneration in Alzheimer’s disease. CNS Neurol. Disord. Drug Targets 13, 1483–4 (2014).

155. Garcia-Esparcia, P. et al. Altered machinery of protein synthesis is region- and stage-dependent and is associated with α-synuclein oligomers in Parkinson’s disease. Acta Neuropathol. Commun. 3, 76 (2015).

156. Uechi, T. et al. Ribosomal Protein Gene Knockdown Causes Developmental Defects in Zebrafish. PLoS One 1, e37 (2006).

157. Crocker, A., Guan, X.-J., Murphy, C. T. & Murthy, M. Cell-Type-Specific Transcriptome Analysis in the Drosophila Mushroom Body Reveals Memory-Related Changes in Gene Expression. Cell Rep. 15, 1580–1596 (2016).

158. G, W., Y, Z., B, C. & J, C. Preliminary studies on Alzheimer’s disease using cDNA microarrays. Mech. Ageing Dev. 124, 115–124 (2003).

159. Taguchi, K. et al. Identification of hippocampus-related candidate genes for Alzheimer’s disease. Ann. Neurol. 57, 585–588 (2005).

160. Martin, I. et al. Ribosomal protein s15 phosphorylation mediates LRRK2 neurodegeneration in Parkinson’s disease. Cell 157, 472–485 (2014).

161. Arnsten, A., Cai, J., Steere, J. & Goldman-Rakic, P. Dopamine D2 receptor mechanisms contribute to age-related cognitive decline: the effects of quinpirole on memory and motor performance in monkeys. J. Neurosci. 15, (1995).

162. Mehta, M. A., Hinton, E. C., Montgomery, A. J., Bantick, R. A. & Grasby, P. M. Sulpiride and mnemonic function: effects of a dopamine D2 receptor antagonist on working memory, emotional memory and long-term memory in healthy volunteers. J. Psychopharmacol. 19, 29–38 (2005).

163. Lee, J. et al. Shank3-mutant mice lacking exon 9 show altered excitation/inhibition balance, enhanced rearing, and spatial memory deficit. Front. Cell. Neurosci. 9, 94 (2015).

164. Mei, Y. et al. Adult restoration of Shank3 expression rescues selective autistic-like phenotypes. Nature 530, 481–484 (2016).

165. Feldmann, R., Denecke, J., Grenzebach, M., Schuierer, G. & Weglage, J. Neurofibromatosis type 1: motor and cognitive function and T2-weighted MRI hyperintensities. Neurology 61, 1725–8 (2003).

166. Picciotto, M. R. et al. Abnormal avoidance learning in mice lacking functional high-affinity nicotine receptor in the brain. Nature 374, 65–67 (1995).

167. Bertrand, D. et al. The CHRNB2 mutation I312M is associated with epilepsy and distinct memory deficits. Neurobiol. Dis. 20, 799–804 (2005).

168. Yang, S. et al. Biochemical, molecular and behavioral phenotypes of Rab3A mutations in the mouse. Genes. Brain. Behav. 6, 77 (2007).

169. D’Adamo, P. et al. Mice deficient for the synaptic vesicle protein Rab3a show impaired spatial reversal learning and increased explorative activity but none of the behavioral changes shown by mice deficient for the Rab3a regulator Gdi1. Eur. J. Neurosci. 19, 1895–1905 (2004).

170. Urigüen, L. et al. Behavioral, neurochemical and morphological changes induced by the overexpression of munc18-1a in brain of mice: relevance to schizophrenia. Transl. Psychiatry 3, e221 (2013).

171. Shapshak, P. et al. Alzheimer’s disease and HIV associated dementia related genes: I. location and function. Bioinformation 2, 348–57 (2008).

172. Baumgärtel, K. et al. Changes in the Proteome after Neuronal *Zif268* Overexpression. J. Proteome Res. 8, 3298–3316 (2009).

173. Smith, A. M., Bowers, B. J., Radcliffe, R. A. & Wehner, J. M. Microarray analysis of the effects of a gamma-protein kinase C null mutation on gene expression in striatum: a role for transthyretin in mutant phenotypes. Behav. Genet. 36, 869–881 (2006).

174. Smith, A. M., Bowers, B. J., Radcliffe, R. A. & Wehner, J. M. Microarray Analysis of the Effects of a γ-protein Kinase C Null Mutation on Gene Expression in Striatum: A Role for Transthyretin in Mutant Phenotypes. Behav. Genet. 36, 869–881 (2006).

175. Davidson, M. E., Kerepesi, L. A., Soto, A. & Chan, V. T. d-Serine exposure resulted in gene expression changes implicated in neurodegenerative disorders and neuronal dysfunction in male Fischer 344 rats. Arch. Toxicol. 83, 747–762 (2009).

176. Espuny-Camacho, I. et al. Hallmarks of Alzheimer’s Disease in Stem-Cell-Derived Human Neurons Transplanted into Mouse Brain. Neuron 93, 1066–1081.e8 (2017).

177. Bado, P. et al. Effects of low-dose d-serine on recognition and working memory in mice. Psychopharmacology (Berl). 218, 461–470 (2011).

178. Barker-Haliski, M. L., Pastuzyn, E. D. & Keefe, K. A. Expression of the core exon-junction complex factor eIF4A3 is increased during spatial exploration and striatally-mediated learning. Neuroscience 226, 51 (2012).

